# The sensitivity of ECG contamination to surgical implantation site in adaptive neurostimulation

**DOI:** 10.1101/2021.01.15.426827

**Authors:** Wolf-Julian Neumann, Majid Memarian Sorkhabi, Moaad Benjaber, Lucia K. Feldmann, Assel Saryyeva, Joachim K. Krauss, Maria Fiorella Contarino, Tomas Sieger, Robert Jech, Gerd Tinkhauser, Claudio Pollo, Chiara Palmisano, Ioannis U. Isaias, Daniel Cummins, Simon J. Little, Philip A. Starr, Vasileios Kokkinos, Schneider Gerd-Helge, Todd Herrington, Peter Brown, R. Mark Richardson, Andrea A. Kühn, Timothy Denison

## Abstract

**Background:** Brain sensing devices are approved today for Parkinson’s, essential tremor, and epilepsy therapies. Clinical decisions for implants are often influenced by the premise that patients will benefit from using sensing technology. However, artifacts, such as ECG contamination, can render such treatments unreliable. Therefore, clinicians need to understand how surgical decisions may affect artifact probability.

**Objectives:** Investigate neural signal contamination with ECG activity in sensing enabled neurostimulation systems, and in particular clinical choices such as implant location that impact signal fidelity.

**Methods:** Electric field modelling and empirical signals from 85 patients were used to investigate the relationship between implant location and ECG contamination.a

**Results:** The impact on neural recordings depends on the difference between ECG signal and noise floor of the electrophysiological recording. Empirically, we demonstrate that severe ECG contamination was more than 3.2x higher in left-sided subclavicular implants (48.3%), when compared to right-sided implants (15.3%). Cranial implants did not show ECG contamination.

**Conclusions:** Given the relative frequency of corrupted neural signals, we conclude that implant location will impact the ability of brain sensing devices to be used for “closed-loop” algorithms. Clinical adjustments such as implant location can significantly affect signal integrity and need consideration.

**Highlights:**

- Chronic embedded brain sensing promises algorithm-based neurostimulation
- Algorithms for closed-loop stimulation can be impaired by artifacts
- The relationship of implant location to cardiac dipole has relevant impact on neural signal fidelity; simple models can provide guidance on the sensitivity
- ECG artifacts are present in up to 50% of neural signals from left subclavicular DBS systems
- Implanting DBS in a right subclavicular location significantly reduces frequency of ECG artifacts
- Cranial-mounted implants are relatively immune to artifacts

## Introduction

Invasive neurostimulation can modulate neural activity and alleviate symptoms in a variety of severe neurological and psychiatric disorders. [1,2] Current advances in deep brain stimulation (DBS) research demonstrate the utility of closed-loop adaptive DBS based on neural feedback signals recorded directly from the stimulation electrodes. [3–6] Most prominently, subthalamic beta activity (13 - 35 Hz) in Parkinson’s disease was shown to reflect parkinsonian motor sign severity [7] that rapidly follows the clinical response to treatment [8–10] and is a promising candidate for adaptive deep brain stimulation (aDBS) [11,12]. Similarly, in patients with epilepsy, seizure activity can inform rapid therapeutic intervention [13]. In such scenarios, clinical success of demand-dependent therapy adaptation depends on the reliability of the feedback signal [14,15]. Neurophysiological recordings are prone to electrical artifacts. The strongest source of electrical activity in the human body is the heart, and the frequency content of cardiac activity overlaps many brain signals of interest (Figure 1). Since the first experience with sensing enabled implantable DBS devices, ECG contamination remains an unresolved problem rendering a significant number of recordings unusable [16,17].Similar issues may arise with motion and muscle contraction. In the present study, we investigate the relationship between the electric field of cardiac activity, implant location, and contamination of neural signals recorded.

**Figure 1:**
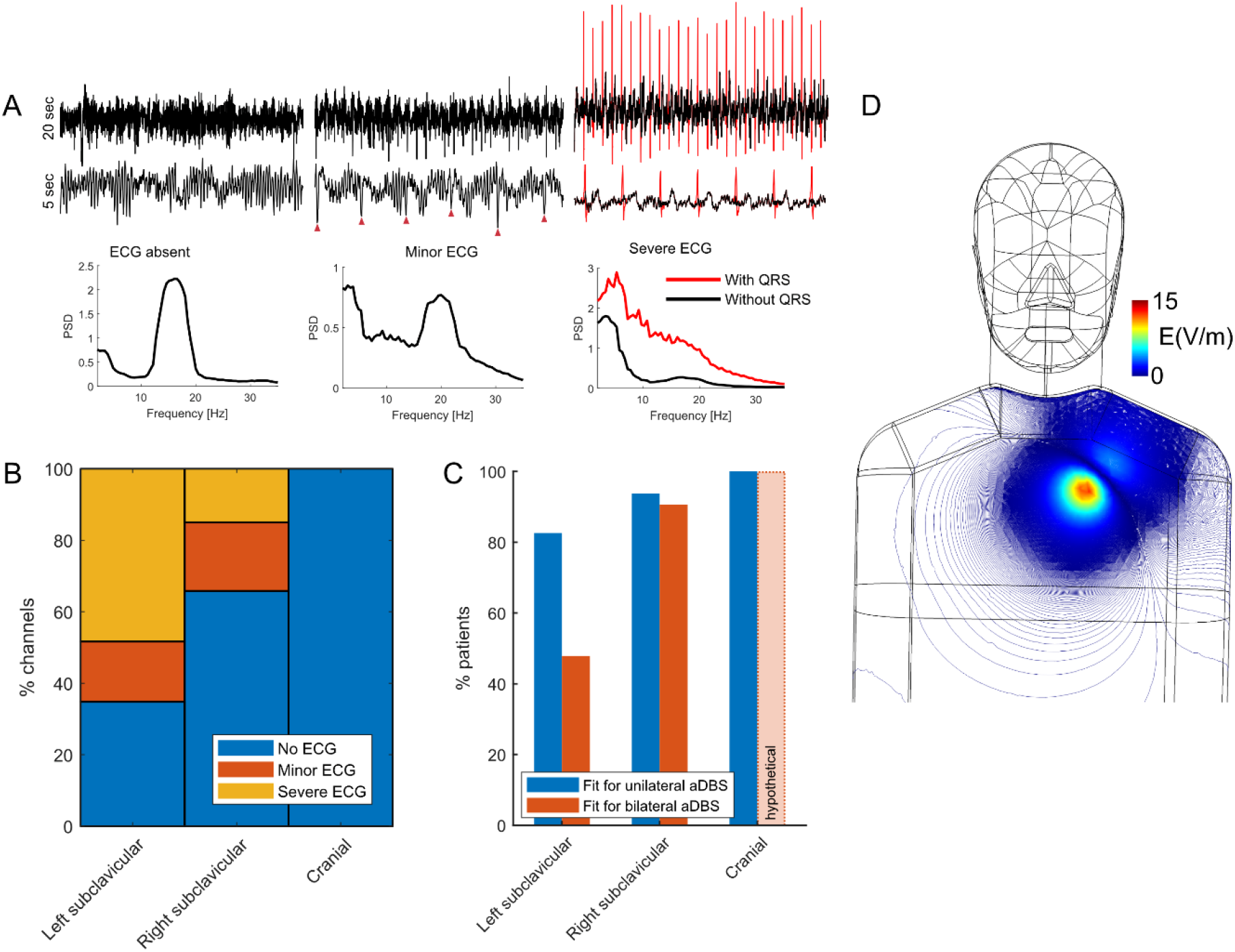
ECG artifacts contaminate neural signals in subclavicular implants. Exemplar subthalamic LFP and resulting power spectral densities (A) recorded from patients with Parkinson’s disease in Berlin demonstrate the artifact categories (absent, minor, severe from left to right). For offline processing, the QRS complex can be identified (e.g. red arrow in minor contamination) and removed (red line in severe contamination category, see https://github.com/neuromodulation/perceive). In the severe contamination example replacing 4.37 s affected by QRS (red high amplitude discharges) with mirrored padding could restore an underlying beta oscillatory peak (black PSD), demonstrating the severity of beta frequency contamination from the QRS complex alone (red PSD). ECG contaminated channels were present in left and right subclavicular implants (B), rendering a significant portion of DBS leads unusable for aDBS trials (C). Modelling the electric field (D) throughout the cardiac cycle suggests a higher susceptibility for ECG artifacts in the left, when compared to right chest.

## Methods

### Empirical data

The study was carried out in accordance with the Declaration of Helsinki and was approved by the internal review board of Charité – Universitätsmedizin Berlin. To validate the predictions of our model, we visually inspected recordings for ECG contamination. Therefore, archival local field potential (LFP) recordings from 8 international neuromodulation centers in 86 implants in 85 patients were inspected for evidence of ECG. From this cohort, 53 patients have undergone implantation of Medtronic Percept DBS pulse generators (21 left subclavicular, 1 left abdomen, 29 right subclavicular, 1 right abdomen, 1 both left and right subclavicular; see Table 1). Most commonly DBS was applied biltaterally (one lead in each hemisphere), 4 implants only had leads in one hemisphere connected (unilateral). Calibration Tests were performed bipolarly for contact pairs 0-2 and 1-3, with 0 being the lowermost contact. Importantly, these calibration tests are performed in passive recharge mode, with stimulation anode and cathode defined as for active stimulation, even when performed at 0 mA. This resulted in 2 channels per lead. One channel from one patient had to be excluded due to impedance issues, resulting in 207 channels overall. Additionally, 32 patients have been implanted with a neurostimulation system mounted to the skull (cranial mount) for treatment refractory epilepsy therapy (RNS Neuropace). For Percept data, bipolar calibration test recordings of ~20 s length were visualized using our open-source Perceive Toolbox (https://www.github.com/neuromodulation/perceive/) in Matlab (The Mathworks). For cranial mounts, visual inspection was performed from routine recordings. ECG artifacts were identified based on the presence of characteristic sharp QRS-like signal deflections of ~150-200 ms width with stereotypic amplitudes occurring at 60-100 bpm. Artifacts were categorized into absent, minor, and severe (see Figure 1), as some recordings had visible but low amplitude contamination, e.g. with just the tip of the QRS complex close to the level of neural activity. Statistical comparison of implant location and observed ECG contamination were performed by aggregating channel counts per implant and using the exact version of non-parametric Wilcoxon’s rank sum tests.

### Modelling

Physiological modeling can be used to better understand the effect of the reference electrode placement on the induced cardiac artifacts [18]. Here, the heart is treated as a single current dipole source in the thorax modelled to have a uniform electrical conductivity [19]. In our computational model using COMSOL software, the current-source dipole heart model is surrounded by a homogeneous volume conductor (average tissue conductivity = 0.33 S/m) with the shape of a three-dimensional human torso, consisting of 2 mm^3^ elements (Figure 1D). The magnitude of the electric current dipole moment is assumed to be 1 (mA meter). This model was solved linearly through finite element methods (FEM) (included in supplemental methods). To predict the maximum possible artifact value, the hypothetical locations of the dipole points are examined in different scenarios around the heart locus. The net voltage induced across the lead was then calculated by integrating the electric field between device location and leads placed in the center of the cranium.

## Results

### Empirical artifact frequency

Visual inspection of LFP recorded in subclavicular implants (N=54 Medtronic Percept, see Figure 1A for exemplar traces) revealed higher proportion of overall ECG contaminated signals (Figure 1B) in left (58/89 channels, 65.2%, N=23 devices) vs. right implants (41/118 channels, 34.8%, N=31 devices, *p*=0.006). Importantly, severe ECG contamination rendering the signals unusable for therapeutic algorithms, were three times more likely to occur in left (43/89, 48.3%, N=23) vs. right implant locations (18/118, 15.3%, N=31, *p*=0.001). Given that each patient has 2 potential recording channels per lead, the availability of at least one usable signal stream per lead and hemisphere is particularly relevant for recruitment and planning of clinical aDBS trials. For bilateral use, at least one unaffected channel per hemisphere and lead is required (Figure 1C). This was the case in only 45.5% (10/22) of patients with left, compared to 89.3% (25/28, *p*=0.002) patients with right implants and two connected leads (unilateral implants excluded). If unilateral recordings were to prove sufficient for bilateral control of the stimulator in aDBS this would have been possible in 63.6% of left implants (14/22) and 96.4% of right implants (27/28, *p*=0.008).

ECG was absent in neural data from cranial implants (32 patients, 128 channels), yielding a significant difference to both left and right subclavicular implants (all *p*<0.01).

### Electric field models predict peak ECG contamination in left chest

The observed differences in cardiac activity are predicted ECG artifact models. The induced voltage at the measurement leads is estimated by integrating the electric fields, approximated using FEM to be 5.02 A/m2 and 15.2 V/m for subclavicular (Figure 1D) and 42.7 μA/m2 and 94 μV/m for cranial implant regions. Cardiac artifacts are estimated to be on the order of 1-3 mV for a chest mounted device and between 4.1 and 6.5 times greater for the left compared to right side, depending on the specific location of the current dipole. The cranial mounted system results in an estimated signal of approximately 100 uV and is less susceptible to specific placement.

This voltage is presented as a common-mode signal to the pre-amplifier. The worst-case susceptibility arises during passive recharge, since this state creates a direct connection to the implantable case, and presents the ECG artifact to the input chain. The final value which is seen in the signal chain depends on the actual common mode rejection ratio (CMRR), which ranges typically typically −60-to-40dB based on matching characteristics of the electrode-lead-extension-generator pathway [20]. Based on this, artifact amplitude can be predicted to exceed the LFP (~1-20 μVrms) in chest-mounted devices. In cranial mount devices the artifact will be below the amplifier noise floor and undetectable.

## Discussion

The present study demonstrates that ECG contamination of neural signals recorded with novel implantable devices attenuates with relation to the electric field of the heart. We derive three major consequences from this. First, for subclavicular systems, the device is highly susceptible to its location relative to the cardiac dipole, if sensing is to be combined with stimulation. In practice, patients with left implants are more likely to exhibit ECG in neural recordings than patients with right implants. From our empirical data, only 45% of patients with left hemibody implants were fit for bilateral adaptive stimulation, compared to 89% in patients with right implants. The second implication is that even right subclavicular implants can suffer from ECG contamination, especially in low amplitude subcortical signals that are being explored for DBS therapies and while applying passive recharge for stimulation; passive recharge is desired to minimize power. The third implication is that, clinical observations made with the IPG uncoupled (e.g. in the Percept BrainSense Survey mode) may have limited merit during stimulation, as the likelihood of artifact contamination is increased when the IPG is coupled for stimulation (even if stimulating at low amplitudes).

Beyond aDBS for Parkinson’s disease, electrophysiological biomarkers have been described in dystonia [21,22], essential tremor [23], Tourette’s syndrome [24,25] and other neuropsychiatric disorders [26]. Further technical improvements for artifact suppression is required to offer new therapeutic advances to all DBS patients.

### Origin of Susceptibility to Artifacts in Brain Sensing Interfaces

Local field potentials are measured as a differential signal from the leads implanted in the brain. The LFP signal can range from 1-20 μVrms [20], and the majority of LFP oscillations are in low frequency bands, ranging from 1 Hz to 100 Hz, where artifacts are also present [27]. When a DBS device is implanted, the device case can act, sometimes inadvertently, as the system’s reference. In theory, the ECG artifact would be rejected by the sensing input chain as a common mode signal. In an implantable system, however, the common mode rejection ratio can be undermined by input impedance mismatch. Such mismatch can occur between the tissue-electrode interface and the front-end amplifier, through impedance differences along the lead and extension interfaces and among some more specific filter and capacitor components used as hardware building blocks in DBS systems.

### Mitigation of ECG contamination in neural recordings from implantable devices

Our study suggests that strategic placement distant to the cardiac electric dipole can partially mitigate ECG contamination. We should note there are alternative approaches to address artifact susceptibility. These methods include improving the matching of the signal chain by improving the electrical properties of leads and extensions, lowering the tissue-electrode interface impedance with new coatings, or exploring alternative signals at frequencies outside of the artifact susceptibility. One or more of these might be adopted in future systems. Moreover, given the proximity of the ECG artifact to the LFP noise floor, higher amplitude signals, e.g. recorded with electrocorticography, will not suffer from ECG and may be viable for certain aDBS applications [12,28–31]. Finally, given the characteristics of the high amplitude QRS component, post-hoc processing can restore a significant portion (~80%) underlying neural activity in many contaminated signals; this approach could prove possible in real-time if achieavable with acceptable power consumption [32].

### Limitations

The evaluation of ECG artefact was visual. In the future we hope to validate our findings with objective measurements of ECG contamination relative to adaptive algorithm requirements. Importantly, even though we demonstrate data from a representative sample size, we only included data recorded from a single device for subclavicular (Medtronic Percept) and cranial (RNS Neuropace) implants. However, a previous generation of subclavicular devices (Medtronic PC+S) had corrupted recording streams often excluded them from otherwise valuable studies [16,17]. For cranial implants, higher amplitude signals from cortical recording locations are additionally beneficial, and may represent a bias in the ECG contamination statistic. It is worth noting that although our focus was on ECG artifacts, susceptibility to motion artifacts raises similar issues that could limit algorithms. The data from cranial mount systems would suggest that the minimal physical motion of the lead and device help mitigate these issues as well, but the root cause of these motion artifacts remains under investigation and could include new processes such as triboelectric phenomena.

In conclusion, sensing enabled IPGs for neurostimulation can suffer from ECG contamination that is larger and more frequent in left subclavicular implant locations, when compared to right-sided or cranial implants. Mitigation strategies include adjustment of implant location, alternative higher amplitude signal sources and post-hoc processing. Given the absence of ECG in cranial implants, our data suggest that future bidirectional brain computer interfaces should explore the utility of cranial mounts to avoid ECG contamination altogether.

## Acknowledgements

The authors would like to thank Dr. R. Zutt (Dept. of Neurology Hagaziekenhuis), Dr. N.A. van der Gaag, and Dr. C.F Hoffmann (Dept of Neurosurgery, Hagaziekenhuis) for the clinical care of the patients and help with data acquisition. This study was funded by the Deutsche Forschungsgemeinschaft (DFG, German Research Foundation) – Project-ID 424778381 – TRR 295 to WJN, IUI and AAK, the Bundesministerium für Bildung und Forschung (BMBF, FKZ01GQ1802) to WJN, the Medical Research Council, United Kingdom (MC_UU_12024/1) to PB, the Czech Ministry of Education under grant AZV: NV19-04-00233 and by Charles University under research project Progres Q27 to RJ.

## Authorship contributions

WJN and TD conceptualized the study, performed statistical analysis, and drafted the manuscript. MMS, MB, PB and TD conceptualized and performed the modelling analysis. LKF, AS, JKK, MFC, TS, RJ, GT, CP, IUI, SL, PS, VK, GH, TH, RMR and AAK acquired and analysed electrophysiological data. AAK and TD provided funding for the study. All authors revised and approved the final version of the manuscript.

